# Unlocking the full potential of Oxford Nanopore reads with NOVOLoci

**DOI:** 10.1101/2025.08.08.669243

**Authors:** Nicolas Dierckxsens, Michael J. Mansfield, Charles Plessy, Nicholas M. Luscombe, Joris R. Vermeesch

## Abstract

Long-read sequencing technologies and accompanying algorithms have substantially advanced genome assembly. However, there are ongoing challenges in assembling complex genomic regions, particularly those associated with genetic disorders. Moreover, assemblies of non-model organisms are often limited by the amount of DNA that can be extracted, so reducing the data modalities that can be obtained. Here, we introduce NOVOLoci, a haplotype-aware assembler capable of high-quality targeted and whole-genome assemblies, despite the relatively high error rates of Oxford Nanopore Technologies (ONT) data. By adopting a novel seed-extension approach with iterative conflict resolution, it achieves accurate haplotype phasing, thus overcoming a critical limitation of current graph-based assemblers. Benchmarking shows that NOVOLoci consistently outperforms the four leading assembly tools across five clinically relevant genomic disorder loci by delivering accurately phased assemblies with superior contiguity and completeness, even compared with hybrid assemblers that combine PacBio and ONT sequencing reads (nearly triple the N90 value compared with Verkko hybrid). We demonstrate its broader utility in assembling the genome of a highly heterozygous non-model animal *Oikopleura dioica*, whose small size limits the amount of DNA available from an individual. Using data only from a single ONT flow cell, NOVOLoci assembles a phased diploid genome with high contiguity and a 32-fold increase in N50 values compared with existing methods. NOVOLoci’s distinct algorithmic strategy demonstrates that substantial advances in genome assembly can still be made. This development brings routine clinical diagnosis of genomic disorders and diploid assembly of non-model organisms with ONT reads a step closer to reality.

## Introduction

Genome assembly remains a challenging task despite the advances in sequencing technologies and assembly algorithms, owing to the complexity and heterogeneity of genomic sequences. The impressive task undertaken by the Telomere-to-Telomere (T2T) Consortium to produce the first complete, gapless sequence of a human genome significantly elevated the standard expected of high-quality assemblies.^1^ This was achieved by combining an unprecedented amount of data (620 Gb) from different sequencing technologies (PacBio, ONT, Illumina for Hi-C and trio-binning) and the development of the new assembly algorithm Verkko^2^, which can integrate both highly accurate PacBio HiFi and long erroneous ONT reads.

However, achieving near T2T assemblies encompassing low complexity and repeat-rich regions remains a formidable challenge^3^. Even with the availability of ultralong reads reaching N50s of ∼100 kb^4^, these regions are frequently associated with gaps or fragmented contigs owing to difficulties in spanning repetitive or duplicated sequences. This limitation is particularly pronounced in genomic regions associated with human genetic disorders, where accurate assembly is critical for clinical applications.

The key limitation of the graph-based algorithms that dominate genome assembly approaches lies in their intrinsic design. Originally developed for high-fidelity short reads, they rely on accurate overlaps between sequences containing haplotype-specific SNP patterns to distinguish between haplotypes^5^. Since sequencing errors in Oxford Nanopore (ONT) reads do not follow a random pattern, real SNPs become indistinguishable from errors, leading to the creation of spurious bubbles in the graph^6^. Software such as Flye resolved this issue by aggressively pruning branches in the graph, resulting in fragmented assemblies and merging of haplotypes into a single path^7^. In contrast, the developers of Hifiasm introduced ONT compatibility following the arrival of more accurate sequencing output; with the advent of R10.4.1 flow cells, Hifiasm can now produce diploid assemblies, although significant quality issues remain (see Results).

Furthermore, most existing algorithms prioritize contig and overall assembly length at the cost of accuracy. Such trade-offs have direct consequences in applications such as clinical genomics in which the diagnosis of genetic disorders demands precision^9^. In such cases, there is need for a more targeted approach to genome assembly that can precisely address the unique challenges posed by specific genomic regions.

Many genetic disorders are attributed to genomic structural rearrangements that are frequently initiated by non-allelic homologous recombination between segmental duplications or low copy repeats^10^. These regions are characterized by their rich content of repeat sequences, copy number variations, and structural rearrangements. Despite significant advances in the identification and characterisation of these disorders, a comprehensive understanding of their genomic architecture at the nucleotide level remains incomplete due to the limitations described above^11^. Accurate haplotype-resolved genome assemblies would enable the precise identification of breakpoints and allele-specific structural variations, thereby enhancing our understanding of the underlying disease pathology. From a clinical standpoint, these advances hold great promise for enhancing genotype-phenotype correlations, personalised diagnostics, and more precise genetic counselling. Ultimately, these developments will contribute to enhanced patient management and therapeutic strategies.

Beyond human genomics, there are challenges in creating *de novo* assemblies for diverse non-model organisms. To achieve diploid assemblies, current approaches generally rely on auxiliary data, such as parental sequences for trio-binning and Hi-C for effective scaffolding and phasing^12^. However, the availability of starting material is frequently a limiting factor. Insufficient DNA means that only a single sequencing run can be performed, leading to decreased sequencing depths, shorter reads, and no additional data available to facilitate phasing.

To address this critical gap, we developed NOVOLoci, a fundamentally different assemble approach that overcomes limitations through a haplotype-aware seed-extension algorithm. NOVOLoci achieves unprecedented accuracy in the most challenging genomic regions while using up to 10-fold less data than current methods, delivering complete, gap-free assemblies where existing methods fail. This enables practical application in individualized patient scenarios, representing a meaningful advance toward clinical genomic applications^13^.

Beyond clinical applications, NOVOLoci addresses fundamental assembly challenges across diverse organisms, as exemplified by *Oikopleura dioica*, a marine tunicate that presents multiple constraints common to non-model organisms. Its small size (∼3 mm) severely limits DNA yield (∼120 ng per adult male), meaning that generally only one type of sequencing data can be produced per individual. Previous assemblies of this species progressed from highly fragmented Sanger-based sequences (1,260 contigs, ∼70 Mb)^14^ to chromosome-scale but haplotype-merged assemblies requiring ONT, Illumina, and Hi-C data from multiple individuals^15^, illustrating the ongoing need for improved assembly methods in non-model organisms. The animal was selected for its ready availability and small genome-size that is amenable for testing.

To demonstrate the effectiveness of our approach across diverse genomic contexts, we benchmarked NOVOLoci’s targeted assembly mode against established methods: Flye^7^ (ONT R9.4.1), Verkko^2^ (PacBio-only and PacBio+ONT hybrid), and Hifiasm^8^ (ONT R10.4.1). Our evaluation for targeted assembly focused on five clinically relevant genomic regions known for structural complexity and association complex neurodevelopmental disorders that affect between 1 in 4,000 people^16^ and 1 in 30,000 live births^17^. For whole-genome assembly, we assessed NOVOLoci against Flye and Shasta to assemble the small tunicate *O. dioica* (∼110 Mb diploid genome).

## Results

### Overview of the NOVOLoci algorithm

NOVOLoci introduces a novel genome assembly approach with superior haplotype phasing capabilities compared with traditional graph-based methods. It currently offers two distinct assembly modes: (i) a targeted mode (T-mode), which uses a seed sequence to initiate assembly and extends contigs to cover specific genomic regions; (ii) and a whole-genome mode (WG-mode), designed to assemble complete genomes by using all input reads as seeds.

The core algorithm uses long-read datasets to iteratively extend seed sequences, resolve haplotypes, and establish consensus sequences (Fig 1). Seeds can be extended unidirectionally or bidirectionally depending on the mode and user requirements. It can handle diverse seed inputs, such as a disease target region, or even homologous sequences from related organisms without incorporating mismatches into the assembly. Optimal seeds are >1000 bp in length with minimal duplications and tandem repeats to prevent erroneous location initiation/early termination.

**Fig. 1.**
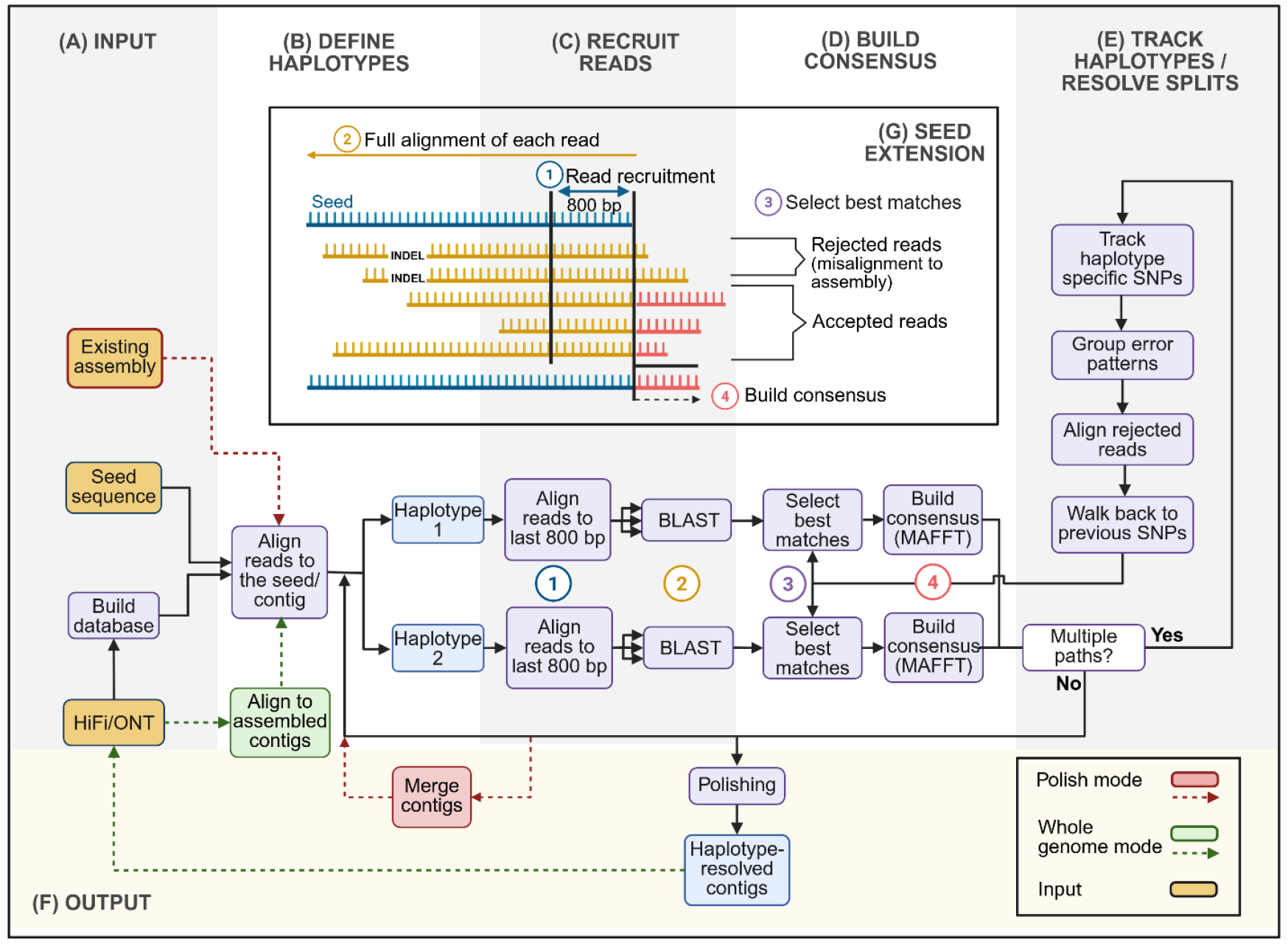
NOVOLoci assembly algorithm. **(A-F)** The NOVOLoci workflow consists of 6 modules, in which modules B to E are iterated until a final assembly is outputted at F. **(G)** Schematic representation of the read recruitment (C) and consensus building steps (D).

NOVOLoci operates through six integrated modules (Fig 1): (A) Input and Database preparation builds a BLAST database from input reads and selects initial seeds; (B) Haplotype detection scans seed sequences for haplotype-specific variation and splits them accordingly; (C) Read recruitment extends seeds by identifying reads mapping to the terminal 800 bp, followed by full alignment to the assembled sequence (both with BLASTn^18^) and acceptance or rejection of reads depending on the completeness of alignment (panel G); (D) Consensus building generates consensus sequences using MAFFT^19^ multiple sequence alignment (panel G); (E) Conflict resolution implements remedial steps when no clear consensus emerges (see Methods); and (F) Assembly finalization terminates the assembly process when either the assembly reaches the requested length (T-mode) or all input reads have been examined for compatibility with seed inputs (WG-mode). Ambiguous nucleotide positions are polished based on the stored consensus values. Potential duplicate contigs are removed, and unique overlapping regions are used to merge contigs. The final output is a separate assembly file for each haplotype and a table with the quality scores for the consensus call of each assembled bp.

### Targeted assembly of complex human genomic regions (T-mode)

Many genomic disorders arise from non-allelic homologous recombination between segmental duplicons, causing recurrent deletions or duplications, and the high variability of these duplicons in size, number, and orientation makes their assembly and mapping especially challenging^20^. We benchmarked the NOVOLoci T-mode against Verkko, Flye, and Hifiasm by assembling five complex disease-associated regions 22q11.2^13^, 15q11-q13^21^, 17p11.2^22^, 16p13.11^23^, and 7q11.23^24^ that are known for their difficulty in assembly. Notably, 22q11.2, 15q11-q13, and 17p11.2 still contain gaps in the GRCh38 human reference genome. We evaluated performance based on the completeness, contiguity, haplotype-phasing, and base-call accuracy, using the T2T Consortium HG002 diploid genome (v1.1) as the gold standard. Each software was tested using default parameters and the appropriate PacBio HiFi and/or ONT R9.4.1 data designed for use as input data (see below for ONT R10.4.1 benchmarks). We used 99 Gb of whole-genome ONT R9.4.1 data which is equivalent of a single PromethION flow cell (average N50 = 84 kb, 15-fold coverage per haplotype) and 96 Gb of PacBio HiFi data.

Overall, NOVOLoci (ONT only) demonstrates the best overall performance across all five genomic regions, consistently achieving the best contiguity (32 contigs total; Fig 2). Flye (ONT), yielded highly fragmented contigs (115 contigs) and was unable to generate phased assemblies, resulting in 21.13 Mb of missing sequence. Verkko using only PacBio data generated phased assemblies of medium completeness (4.44 Mb of missing sequence), but poor contiguity (480 contigs); whereas Verkko’s PacBio+ONT hybrid assemblies displayed improved performance (only 1 Mb of missing sequence and 110 contigs). PacBio-supported assemblies generally exhibited better base-level accuracy owing to the higher accuracy of the sequencing technology itself (QV >40); nonetheless ONT R9.4.1-only assemblies achieve QV scores above 29 (>99.85%), which is more than sufficient for accurate SV detection and structural rearrangement analysis (see below).

**Fig. 2.**
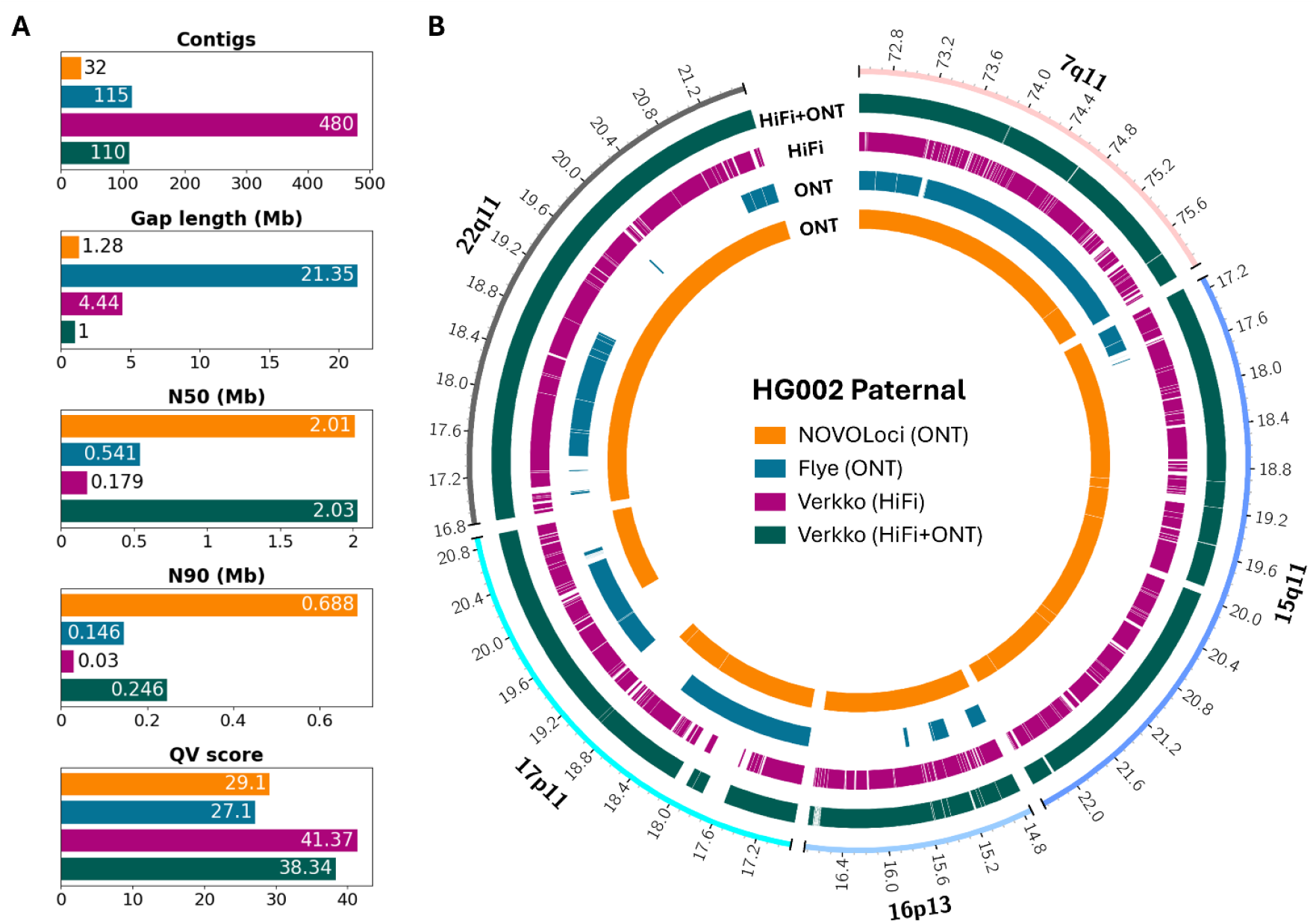
Targeted assembly benchmark for 5 complex regions. (A) Benchmarks statistics for the 5 complex regions of both haplotypes combined. (B) Alignment plot of the paternal assemblies of each of the 5 complex regions against the T2T Consortium HG002 diploid genome (v1.1) as the gold standard.

Below we examined each of the five complex region in detail to understand the specific assembly challenges and NOVOLoci’s performance advantages. We focus comparisons with Verkko’s hybrid (PacBio+ONT) assemblies (Fig 2).

The 7q11.23 region (3.5 Mb) associated with the Williams and 7q11.23 Duplication Syndromes is characterized by three primary segmental duplicons, designated as centromeric, medial, and telomeric. The duplicons vary in size, typically ranging from ∼320 kb to ∼500 kb^25^. NOVOLoci resolved both haplotypes as two overlapping contigs. Verkko hybrid generated 3 contigs with a large 253 kb gap for the maternal haplotype and 5 contigs with 10 kb of gaps for the paternal haplotype.

The 6.5 Mb 15q11–q13 region is characterized by a complex architecture of segmental duplications that are organized into five breakpoint regions, each containing duplicated sequences that vary in size and orientation. These duplications range from ∼200 kb to ∼400 kb in length and share high sequence identities, facilitating misalignments during meiosis and so causing deletions or duplications associated with the Prader-Willi,Angelman and 15q11–q13 Duplication Syndromes^26-28^. NOVOLoci produced fully phased assemblies comprising 7 contigs for both the paternal and maternal haplotype. The maternal haplotype has a gap of 420 kb caused by the merging of two highly homozygous regions between haplotypes. Verkko hybrid did not match this level of contiguity and completeness, with 15 and 10 contigs and 29 kb and 279 kb missing for the maternal and paternal haplotypes, respectively.

The 16p13.11 region (2 Mb) – whose micro-deletions and -duplications are associated with a broad spectrum of neurodevelopmental disorders – is organized into three main intervals (I–III), each containing segmental duplicons that vary in size and orientation. Duplications within these intervals range from ∼0.79 Mb to ∼2.82 Mb in length and share high sequence identity^29^. NOVOLoci achieved single contigs for each haplotype. In contrast, Verkko hybrid outputted 38 and 25 contigs and 19 kb and 26 kb missing respectively for the maternal and paternal haplotypes.

The 17p11.2 region (4 Mb) contains four complex segmental duplicons known as SMS-REPs that vary in size between ∼150 kb and ∼250 kb. These repeats flank the commonly deleted or duplicated segments and are the major drivers of non-allelic homologous recombination events associated with Smith-Magenis and Potocki-Lupski Syndromes^30^. Although NOVOLoci did not fully cover the region, leaving a 854 kb gap in the paternal haplotype due to the merging of two highly homozygous regions between haplotypes, with a total of 9 contigs, it still achieved the most contiguous assembly compared with Verkko hybrid’s total of 12 contigs and 382 kb missing.

The 22q11.2 region harbours four segmental duplicons each containing multiple smaller duplicons varying in size from ∼25 to ∼155 kb^31^. Here we assembled a 5 Mb region encompassing these four critical duplicon blocks linked to 22q11 Deletion Syndrome (formerly including DiGeorge and Velocardiofacial Syndromes). Using ONT R9.4.1 data only, NOVOLoci successfully produced a single contiguous assembly for the paternal haplotype and two overlapping contigs for the maternal haplotype, almost matching the performance of Verkko’s hybrid (PacBio+ONT) method.

These targeted assembly benchmarks collectively highlight NOVOLoci’s substantial improvements in assembling and phasing clinically significant genomic regions, demonstrating its robust capability in accurately reconstructing complex genomic architectures.

### Comparison with Hifiasm using ONT R10.4.1 flow cell data (T-mode)

The latest ONT R10.4.1 flow cell combined with Kit 14 gives a significant improvement in base call accuracy, yielding single molecule reads with QV20+ scores. This prompted the development of a new Hifiasm module that works with ONT data alone. To assess the capabilities of Hifiasm and NOVOLoci, we benchmarked the same five complex regions using publicly available R10.4.1 ONT data (Fig 3). We used 110 Gb of whole-genome R10.4.1 ONT data from two flow cells (average N50 = 109 kb, 17-fold coverage per haplotype).

**Fig. 3.**
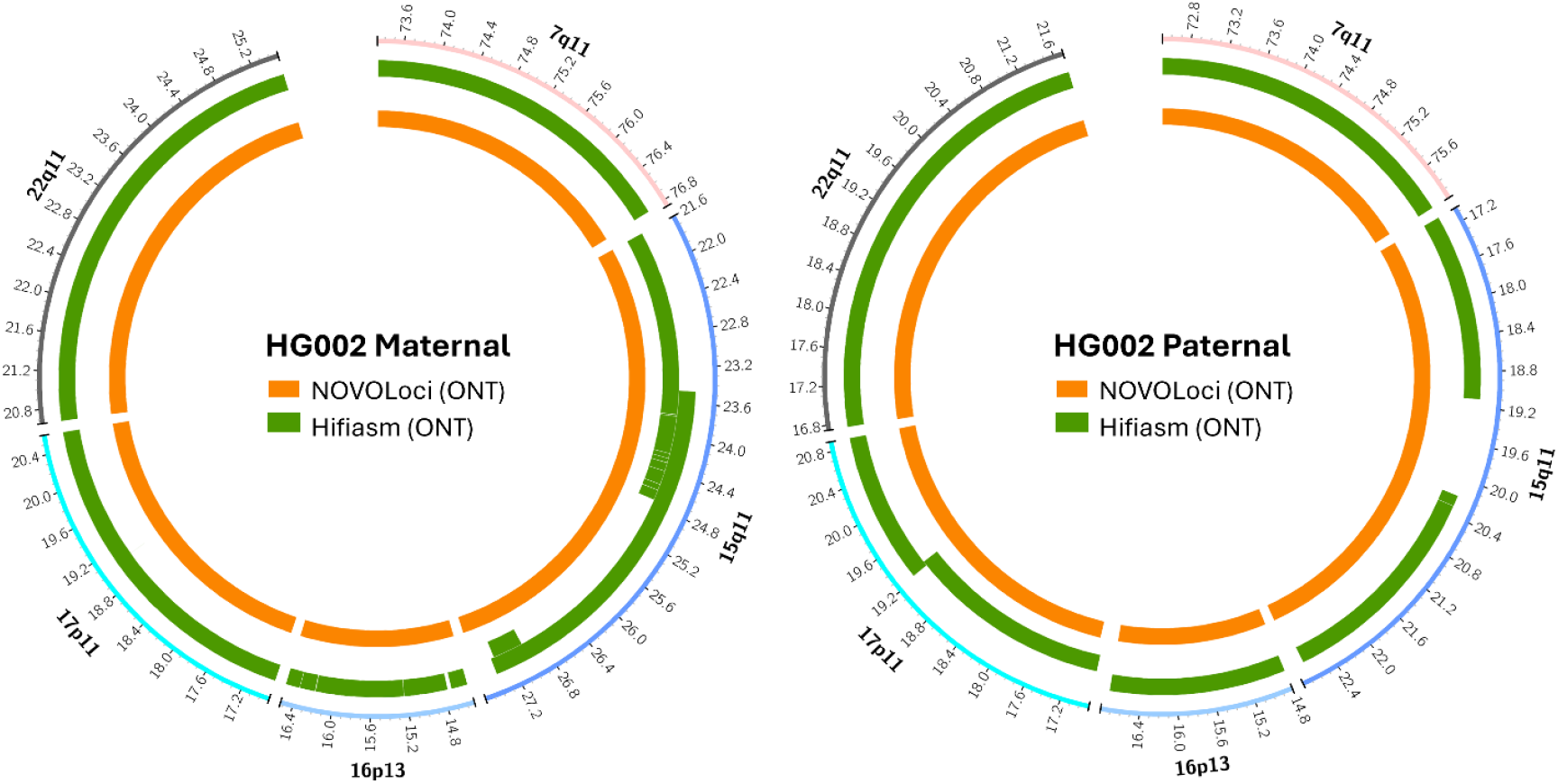
R10.4.1 Nanopore assemblies with NOVOLoci and Hifiasm for 5 complex regions. Alignment plots of the maternal and paternal assemblies of each of the 5 complex regions against the T2T Consortium HG002 diploid genome (v1.1) as the gold standard.

The first major difference we observed compared with R9.4.1 was that both Hifiasm and NOVOLoci required data from two flow cells, rather than one, to produce accurate phased assemblies for each region. This requirement was due to the lower sequencing depth of R10.4.1 flow cells (55 Gb, 8.5-fold coverage per haplotype) compared with R9.4.1 (99 Gb, 15-fold coverage per haplotype).

Compared with R9.4.1 ONT reads, NOVOLoci successfully assembled each of the five regions into a single contig per haplotype. At first glance, Hifiasm appeared to perform only slightly less well than NOVOLoci (14 contigs compared with 10); but, upon alignment to the reference genome, we found that all but 2 contigs were chimeric, consisting of alternating blocks from each parental haplotype. Splitting contigs into correctly phased blocks, yields 36 phased alignments. This suggests that Hifiasm prioritizes contiguity to achieve high N50 values over precise haplotype phasing. This behaviour was not observed with PacBio HiFi data, for which Hifiasm was originally designed.

QV scores for both NOVOLoci and Hifiasm are similar, ranging from 31 to 36, which corresponds to 99.92% - 99.98% nucleotide accuracy. Closer inspection of the assembly errors of both methods revealed that 93% are indels occurring in A/T homopolymers longer than 5 bp.

### Diploid assembly of a small non-model animal genome using only R9.4.1 ONT data (WG-mode)

Next, we tested NOVOLoci’s ability to assemble a complete haplotype-phased genome using DNA extracted from an individual male *O. dioica*, a marine tunicate whose genome presents assembly challenges that are common for non-model organisms. Its small size limits the amount of DNA that can be extracted from an individual (∼120 ng per adult male) and the high heterozygosity can cause difficulties in producing haplotype-merged assemblies. The exceptionally small haploid genome consisting of two autosomes, and sex chromosomes comprising pseudo-autosomal and X/Y-specific regions presents a convenient test-case for benchmarking. We used 9.25 Gb of ONT R9.4.1 data from a single MinION flow cell (average read N50 = 10.7 kb, 84-fold coverage), representing the typical DNA yield limitations encountered with small organisms. No additional sequencing data or scaffolding information was used.

The NOVOLoci WG-mode produced a markedly superior assembly compared with existing software (Fig 4), outputting 27 contigs (N50 = 6.61 Mb) whose total 109 Mb length approached the expected ∼110 Mb diploid genome size. Flye generated a highly fragmented assembly comprising 4,280 contigs (104.0 Mb total, N50 = 0.21 Mb), whereas Shasta produced 923 contigs representing less than ∼70 % of the full genome (73.7 Mb total, N50 = 0.15 Mb). Parameter optimization confirmed that these performance differences were not due to suboptimal settings (Supplementary Tables 1). NOVOLoci assembles most of the 12 chromosome arms (6 haploid chromosomes with short and long arms each) in 2 or 3 contigs. The largest contigs belong to the haploid X-specific region (∼12.7 Mb), which was more easily assembled as it lacked the complexity of diploid chromosomes, and both Shasta and Flye were able to produce contigs spanning most or all the X-specific region after searching for suitable non-default parameters.

**Fig. 4.**
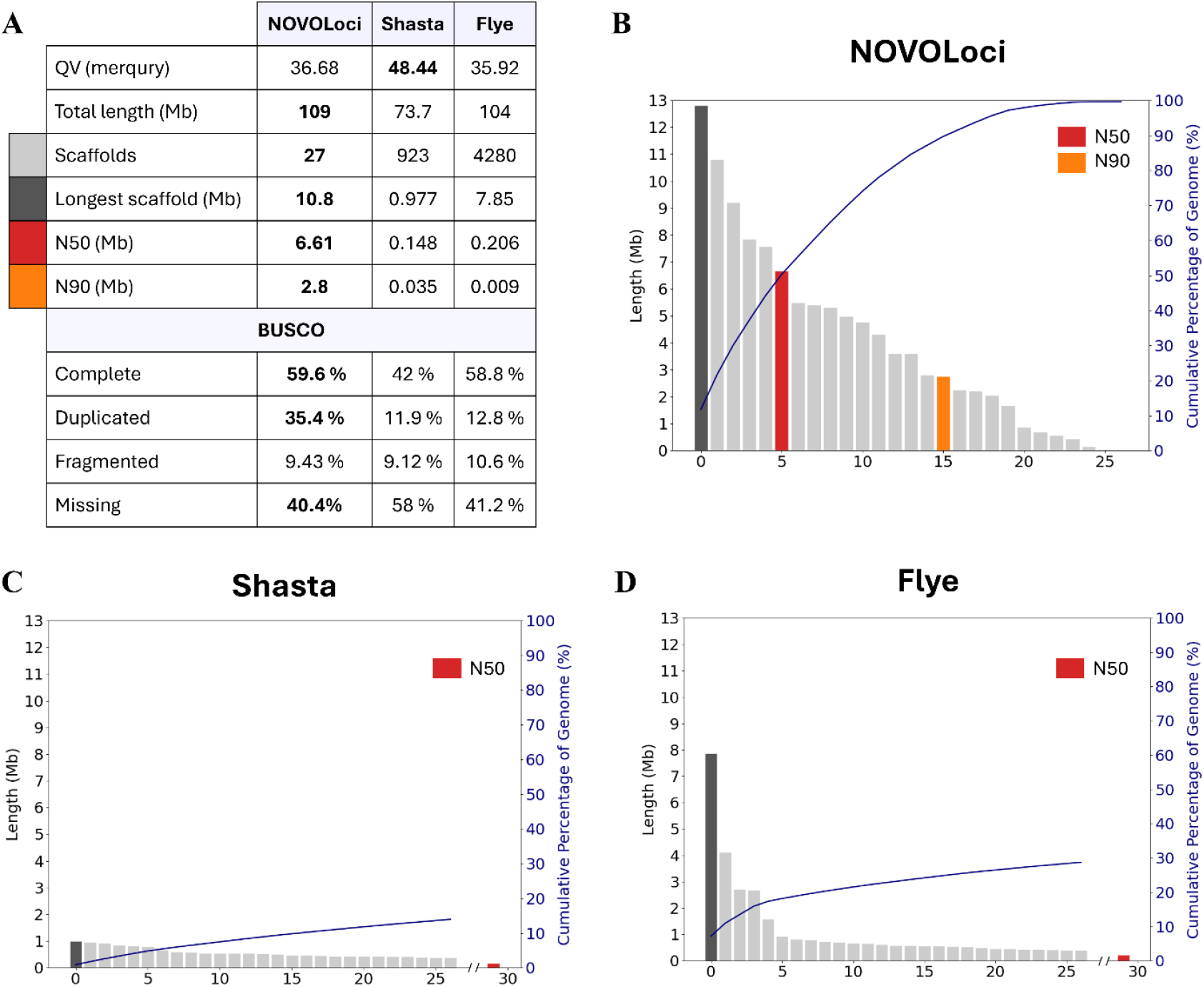
Whole genome benchmark for the assembly of *O. dioica*. (B-D) Visual representation of the lengths of the 27 largest contigs for each assembly. The contig that represents the N50 value is coloured red, which was out of range of the first 27 for Shasta (C) and Flye (D).

NOVOLoci achieved an average QV score of 36.7 (= 99.98 % accuracy), which is comparable to Flye (35.9, 99.97 %). Shasta achieved a higher average QV score (48.4, 99.998 %), as the assembly covered only the most easily produced regions. Gene-level completeness assessment using BUSCO revealed 569 complete genes (59.6% of metazoan OrthoDB set, n=954) for NOVOLoci versus 549 for Flye and 192 for Shasta. The relatively low BUSCO scores reflect non-canonical splice sites, which are not handled well by BUSCO, and extensive known gene losses in *O. dioica*, rather than assembly deficiencies. Importantly, NOVOLoci identified more duplicated BUSCO genes (231 vs 141 for Flye and 28 for Shasta), indicating successful resolution of both haplotypes.

These results demonstrate NOVOLoci’s ability to generate near-complete diploid assemblies from very limited input data, achieving >32-fold improvement in contiguity over existing methods. The successful haplotype phasing represents the first diploid assembly of *O. dioica* and demonstrates NOVOLoci’s utility for non-model organisms where limited DNA availability prevents a comprehensive set of sequencing data to be provided.

## Discussion

NOVOLoci represents a substantial advance in precision genome assembly, especially for highly complex, medically significant genomic regions and organisms with small genomes. By employing a seed-extension algorithm combined with adaptive error resolution strategies, NOVOLoci effectively overcomes the limitations commonly faced by graph-based assemblers.

Accurate haplotype-resolved assemblies can enable the characterization of even the most complex duplicons. This allows for the precise identification of genes located within duplicons, mapping of their variation and that of their paralogues, and the assessment of evolutionary constraint acting on these genes. These advances pave the way for targeted diagnostic screening of pathogenic variants within duplicated regions. The benchmarks on duplicon-rich regions demonstrate NOVOLoci’s ability to generate accurate and contiguous assemblies. NOVOLoci’s ONT-only assemblies outperformed all alternative methods, including Verkko’s hybrid (HiFi+ONT) approach.

The increased base call quality with the latest R10.4.1 flow cells make it possible for Hifiasm to attempt phased assemblies with ONT data only. However, although haplotypes with impressive N50 values are outputted, all but two are chimeric (mixing the two haplotypes). This appears to arise from the prioritization of contiguity over precise phasing. For 15q11 and 16p13 the switching arose within the complex region, whereas the others occurred at loci flanking the target region. This is problematic nonetheless, since in the absence of a reference genome, the widespread occurrence of chimeric contigs makes it impossible to detect the errors. Both evolutionary analyses and clinically meaningful variant interpretation demand accurate, haplotype-resolved genome assemblies. Chimeric contigs that merge sequences from both haplotypes can obscure allelic relationships, mask compound heterozygosity, and falsely suggest or conceal gene dosage imbalances. This compromises the ability to distinguish between dominant and recessive inheritance patterns, a critical requirement for identifying novel disease-associated genes and correctly interpreting the pathogenicity of variants. Therefore, avoiding chimeric assemblies is essential to enable reliable diagnostic interpretation and gene discovery in duplicated regions of the genome. NOVOLoci assembled all complex haplotypes as individual contigs without any phasing errors.

The chimeric assembly problem and the inability to resolve haplotypes extends beyond clinical applications to non-model organism genomics, where reference genomes are unavailable. Many non-model organism projects face additional constraints: limited DNA availability, high heterozygosity, and inability to generate multiple data types from single individuals. These challenges often force researchers to accept haplotype-merged assemblies that obscure important biological variation. As demonstrated by *O. dioica*, NOVOLoci addresses these fundamental limitations, making it particularly valuable for projects where DNA scarcity restricts sequencing options or where resolving haplotype-specific variation is critical for understanding evolutionary or ecological processes.

A current limitation of the updated ONT R10.4.1 technology is the reduced output, meaning that two flow cells are now required to achieve sufficient sequencing depth for successful assembly with either NOVOLoci or Hifiasm. This limits ONT’s applicability for clinical diagnostic purposes. We eagerly await future improvements to the yield of R10.4.1 flow cells or the release of new clinical use flow cells that strike a balance between the higher yield of R9.4.1 and the improved base-calling accuracy of R10.4.1.

An important consideration in choosing an assembly method is the trade-off between accuracy and computational speed. NOVOLoci’s intense focus on achieving highly accurate, phased assemblies results in considerably slower assembly times compared with graph-based assemblers: for instance, the 110 Mb *O. dioica* genome required a maximum 5 GB of RAM - easily within the capacity of most desktop computers - but took 111 hours to complete (compared with 2.6 hours with a max of 57 GB RAM for Flye and 1 hour with a max of 50 GB RAM for Shasta). As the run-time scales linearly with assembly size (target region or whole genome), meaning it would be impractical to attempt assembling most large genomes. Other factors that affect speed are sequence complexity, the presence of homozygous stretches, read quality and sequencing depth.

Therefore, in practice, we currently recommend limiting the assembly size to regions <20 Mb in T-mode and diploid genomes that are <250 Mb in WG-mode, with a minimum sequencing depth of 10x per haplotype. This places NOVOLoci as a precision tool, in cases where accuracy, phasing and data availability are paramount considerations.

In the near future, to take advantage of the speed of graph-based assemblies and the high accuracy afforded by NOVOLoci, we shall implement a polish mode (P-mode), which completes gaps, corrects phasing and merges contigs outputted by other assemblers to refine and enhance the final assembly. Moreover, since ∼70% of the runtime involves the multiple sequence alignment step using MAFFT, we are developing a new consensus-calling module that replaces MAFFT altogether, which should at least double the compute speed. The main hurdle for this polishing mode is obtaining an existing assembly that contains a minimal of phasing errors, which are prevalent with current graph-based assembly algorithms. NOVOLoci currently exclusively supports ONT reads; therefore, in addition to the above to speed improvements, we are currently implementing support for PacBio reads and hybrid datasets. Further updates will expand beyond haploid and diploid genomes, to encompass polyploid genomes common in plants, and genome-assembly from metagenomic data. The success of NOVOLoci’s seed-extension approach suggests broader applications for iterative consensus-building algorithms in challenging genomic tasks and these planned updates will expand NOVOLoci’s versatility and potential application scope.

## Methods

NOVOLoci is a seed-extend based assembler based on the same principle as the organelle assembler NOVOPlasty^32,33^, with the difference that it is built for long sequencing reads and includes BLAST and MAFFT as a backbone of the algorithm. It starts with storing the sequences into a local BLAST database, which allows quick accessibility of the reads (Fig 1).

### Read recruitment with BLAST

The basic principle of the assembly process is to first identify putative matches by aligning the last 800 bp of the assembly to all the reads in the database, followed by a full BLASTn alignment of the selected reads to the complete assembly. All mismatch positions between the aligned reads and the assembly are stored to resolve potential conflicts during consensus building. The aligned reads are ranked by the relative cumulative score of the alignment length, accuracy and the amount of verified haplotype-specific SNPs. The best scoring matches (max. 30) are used to build a consensus for each succedent 800 bp of the extension by generating multiple alignment files with MAFFT.

### Complex regions resolution

In case of conflicts during the consensus, which can arise at the end of duplicated or homozygous regions, several modules are activated to resolve the assembly (Fig 5).

**Fig. 5.**
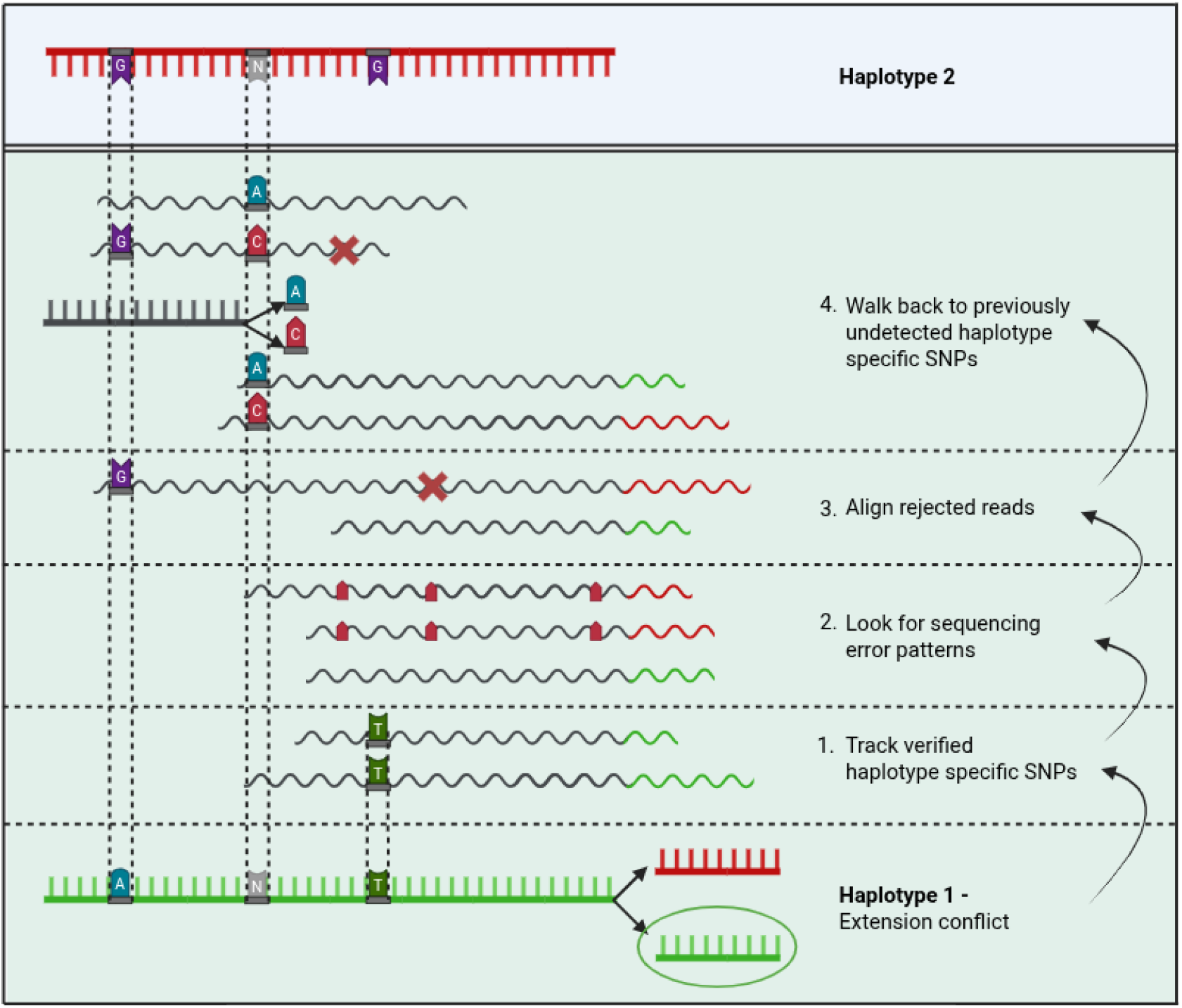
Consensus conflict resolution modules. NOVOLoci will initiate several modules (1–4) step by step to resolve conflicts during consensus calling, which arise from differences in the other haplotype or from duplicated regions.

First, extensions are filtered based on previously confirmed haplotype-specific SNPs, followed by matching error patterns from the individual read alignments with the consensus patterns. When still unresolved, the previous steps are repeated after rejected reads which are longer than the longest read from either of the consensus read groups are included in the consensus call. As a final step, NOVOLoci will look for previously unresolved SNP patterns in the aligned reads that match the consensus patterns and restart the assembly at one of these positions with the knowledge that this position is a true SNP and should be resolved. Recursively jumping back to previously undetected true SNP positions makes it possible to phase long homozygous regions that differ by only a handful of SNPs. When there is still more than one possible consensus extension and it is not possible to resolve, the assembly will be terminated or new seeds are generated from each consensus pattern.

### DNA extraction and sequencing

*O. dioica* was collected from Kin Bay in Okinawa Prefecture, Japan, and cultured in the laboratory at OIST^15^. genomic DNA was extracted using a modified salting-out protocol^15^, processed using the Ligation Sequencing Kit (Nanopore LSK109) and sequenced using a single ONT R9.4.1 Minion flow cell.

### Targeted assembly benchmark

Public Nanopore data was obtained from the Human Pangenome Project database^34^. Nanopore sequencing was performed with the ultralong protocol and re-base-called with Guppy v6.3.7. We selected one dataset originating from a single flow cell, with an average N50 of 84 kb and a sequencing depth of 30x. This data was used for the NOVOLoci (v0.3), Flye (v2.9.1) and hybrid Verkko (v2.2.1) assemblies. For PacBio data, we chose to retain sequencing data and the benchmark assemblies from the Verkko manuscript^20^. Both were downloaded from respectively the Verkko GitHub page (https://github.com/marbl/verkko/blob/master/paper) and Zenodo (https://zenodo.org/records/7400747). The PacBio sequencing data was used to run the hybrid assembly with Verkko. For the ONT R10.4.1 benchmark against Hifiasm (v0.24.0), we selected 2 ONT runs with an average N50 of 109 kb and a combined sequencing depth of 34x. Links to all datasets used in this study are provided in the Data Availability section. Version 1.1 (July, 2024) of HG002 was used as a reference genome and was downloaded from https://github.com/marbl/hg002. The five complex regions were extracted from the whole genome assemblies by aligning them against the HG002 reference genome with minimap2 and extracting the contigs that overlap the coordinates of each of the five regions. Since minimap2 has a high tolerance for miss assemblies or incorrect phasing, we included an additional step that splits alignments when there is a deletion or insertion above 50 bp in the assembly compared to the reference. All seed sequences and configuration files used for NOVOLoci assemblies can be found on its GitHub page: https://github.com/ndierckx/NOVOLoci/tree/master/Benchmark

### Whole genome assembly benchmark

Whole-genome assemblies for an individual male *O. dioica* were produced with Flye (v2.9.1), Shasta (v0.11.1), and NOVOloci (v0.2). Assembly statistics are tabulated in Figure 3. Default parameters were used to generate the Flye and Shasta (using the Nanopore-Phased-May2022 config) assemblies. However, several input parameters were assessed to ensure that any difficulties in assembly were not due a particularly unfavourable set of default parameters. For Flye, the – nano-corr and –nano-hq accuracy presets were assessed with and without the –keep-haplotypes flag. For Shasta, different config files (corresponding to different accuracy presets) with different settings for minReadLen, detangleMethod, assemblyMode, and markerGraphLowCoverageThreshold. Per-base accuracy was assessed with merqury (version 1.3) as well as BUSCO (version 5.5.0) using the OrthoDB metazoa database (version 10, 954 BUSCO genes). All seed sequences and configuration files used for NOVOLoci assemblies can be found on its GitHub page: https://github.com/ndierckx/NOVOLoci/tree/master/Benchmark

## Conclusion

NOVOLoci significantly advances genome assembly by providing highly accurate, contiguous, and haplotype-resolved assemblies, particularly from complex genomic regions using solely error-prone long-read sequencing data. Its unique capacity for accurate phasing of diploid genomes positions it distinctly within the current methodological landscape, addressing critical gaps left by traditional assembly approaches. Although computationally intensive, ongoing optimizations promise greater speed and broader utility, making NOVOLoci increasingly accessible for diverse genomic research contexts. By facilitating precision assemblies from limited input data, NOVOLoci opens promising avenues for genomic research and clinical diagnostics, underscoring its transformative potential for understanding genetic disorders, biodiversity, and genome evolution.

## Declarations

N.D. is a recipient of the OIST Interdisciplinary Postdoctoral Scholar Fellowship (IPSF) and the Post Doctoral Mandate (PDM) of KU Leuven. The research was funded by Fonds Wetenschappelijk Onderzoek (grant numbers G0E1117N and G0A2622N to J.R.V.), KU Leuven (C14/22/125 to J.R.V.), and core-funding to the Genomics & Regulatory Systems Unit from OIST.

## Acknowledgements

We thank the Scientific Computing and Data Analysis Section of the Research Support Division at OIST for their support.

## Data availability

All human HG002 ONT datasets were obtained from the Human Pangenome Research Consortium (HPRC). The R9.4.1.1 ONT dataset: https://s3-us-west-2.amazonaws.com/human-pangenomics/NHGRI_UCSC_panel/HG002/nanopore/ultra-long/03_08_22_R941_HG002_rebasecalling-guppy-6.3.7/03_08_22_R941_HG002_1.fq.gz The R10.4.1 ONT datasets: https://s3-us-west-2.amazonaws.com/human-pangenomics/submissions/5b73fa0e-658a-4248-b2b8-cd16155bc157--UCSC_GIAB_R1041_nanopore/HG002_R1041_UL/Guppy6/HG002_1_R1041_ULCIR_Guppy_6.3.7_5mc_cg_sup_prom_pass.fastq.gz and https://s3-us-west-2.amazonaws.com/human-pangenomics/submissions/5b73fa0e-658a-4248-b2b8-cd16155bc157--UCSC_GIAB_R1041_nanopore/HG002_R1041_UL/Guppy6/HG002_2_R1041_ULCIR_Guppy_6.3.7_5mc_cg_sup_prom_pass.fastq.gz

For the HG002 human reference genome, we used hg002v1.1from the T2T consortium: https://s3-us-west-2.amazonaws.com/human-pangenomics/T2T/HG002/assemblies/hg002v1.1.fasta.gz.

The ONT reads from Oikopleura can be found in the European Nucleotide Archive (ENA) under accession number ERR15172616. All the assemblies and seed sequences used for the targeted assembly with NOVOLoci can be found on the github page: https://github.com/ndierckx/NOVOLoci

## Author contributions

Conceptualization: N.D.

Methodology: N.D.

Development: N.D.

Formal analysis: N.D., M.J.M., C.P.

Writing - original draft: N.D., M.J.M., N.M.L.

Writing - review & editing: C.P., J.V.

Resources: N.D., N.M.L., J.V.

Supervision: N.M.L., J.V.

